# The Establishment of Prostate-specific, SKP2 Humanized Mice by CRISPR Knock-in Method Reveals Neoplastic Initiation and Microenvironmental Reprogramming

**DOI:** 10.1101/2025.05.09.651611

**Authors:** Liankun Song, Vyvyan Nguyen, Shan Xu, Kelly Vy Tuong Ho, Bang H. Hoang, Jianhua Yu, Edward Uchio, Xiaolin Zi

## Abstract

A recent study has shown that SKP2 inactivation can prevent cancer initiation by extension of total cell cycle duration without perturbing normal division, which suggests a new strategy for cancer prevention. However, direct in vivo evidence for human SKP2 on cancer initiation and prostatic microenvironment is still lacking and a prostate-specific SKP2 humanized mouse model is critical for developing prostate cancer immunoprevention approaches through targeting human SKP2. We therefore have established a prostate-specific human *SKP2* (h*SKP2*) knock-in mouse model by a CRISPR knock-in approach. Overexpression of h*SKP2,* which is driven by an endogenous mouse probasin promoter, induces prostatic lesions including hyperplasia, mouse prostate intraepithelial neoplasia (mPIN), and low-grade carcinoma and increases prostate weights. Transcriptional profiling by RNA-sequencing analysis revealed significant gene expression alterations in epithelial to mesenchymal transition (EMT), extracellular matrix, and interferon signaling in the prostate of h*SKP2* knock-in mice compared to wild-type mice. Single cell deconvolution showed an increase of fibroblasts population and a decrease of CD8^+^ T cell and B cell populations in the prostate of *hSKP2*-knock-in mice. Consistently with these results from the SKP2 humanized mouse, overexpression of *hSKP2* in human prostate cancer PC3 cells markedly increased cell migration and invasion and induced the gene expression of EMT and interferon pathways, including *FMOD, THY1, PFKP, USP18, IL15,* etc. In addition, paired prostate organoids were derived from SKP2 humanized and wild-type mice for drug screening and validated by known SKP2 inhibitors, Flavokawain A and C1. Both of which selectively decrease the viability and alter the morphologies of organoids of h*SKP2* knock-in rather than wild-type mice. Our studies provide a well-characterized prostate-specific h*SKP2* knock-in mouse model and offer new mechanistic insights for understanding the oncogenic role of SKP2 in shaping the prostatic microenvironment during early carcinogenesis.

## Introduction

S-phase kinase associated protein 2 (SKP2) is the rate-limiting component of the Skp, Cullin, and F-box (SCF) containing E3 ubiquitin ligase complex that catalyzes the ubiquitination of proteins for targeted degradation by 26S proteasome [1–3]. SKP2 plays a central role in cell cycle regulation, cellular senescence, and apoptosis by targeting and ubiquitinating cell-cycle and apoptosis regulators, including E2F1, the Forkhead box O (FOXO) family members. SKP2 binds and degrades cyclin-dependent kinase inhibitors such as p27^Kip1^ and p21 for controlling cell cycle entry and G1/S transition and then cell proliferation [1–3].

SKP2 has been shown to function as a proto-oncogene causing the pathogenesis of lymphomas [4]. Overexpression of SKP2 is frequently observed and associated with cancer progression, metastasis, and poor prognosis in a wide variety of cancers including prostate cancer [4, 5]. In clinical studies of prostate cancer, SKP2 has been reported to be a key amplified gene [6]. SKP2 overexpression has been found in 86.4 % (64 of 74 samples) of pre-malignant high grade-prostatic intraepithelial neoplasia (HG-PIN) and in 557 of 622 (89.5%) primary prostate cancer specimens, showing a strong positive association with pre-operative serum prostate specific antigen (PSA) levels, Gleason score, tumor grade, and biochemical failure in men treated by prostatectomy [7–11]. There are numerous studies using a *Skp2* knockout mouse model. These studies demonstrate that *Skp2* deficiency in mouse models abolishes or inhibits spontaneous tumorigenesis that were initiated by inactivation of PTEN, ARF, pRB as well as by overexpression of Her2, through inducing p53-independent cellular senescence or blocking Akt-mediated aerobic glycolysis [12–15]. A very recent study published in Nature by Chen et al. [16] reported that cells with shorter cell cycle durations are more susceptible to oncogenic transformation than those with longer cell cycles and genetic inactivation of Skp2, a key determinant of cell cycle duration, completely block tumorigenesis in mice. These results suggest a previously unrecognized mechanism for developing new cancer preventive or therapeutic strategies through targeting SKP2.

However, compared to large amounts of studies using a *Skp2* knockout mouse model, direct in vivo evidence for human SKP2 on tumor initiation and prostatic microenvironment is still lacking. In addition, to develop prostate cancer immunoprevention and immunotherapy approaches by targeting human SKP2 oncogene and its associate pathways, a prostate-specific SKP2 humanized mouse model is critical. We therefore have established a more pathophysiologically relevant prostate carcinogenesis model by knocking-in of human *SKP2* gene into a locus that is regulated by the endogenous *probasin* promoter using the CRISPR/Cas9 method. We have also characterized molecular and cell population alterations in the prostate of human *SKP2* (h*SKP2*-KI) mice. The results demonstrate a profound remodeling of the prostatic microenvironment through h*SKP2* overexpression-induced dysregulation of EMT and interferon pathways. Single cell deconvolution shows an increase of fibroblasts population and a decrease of CD8^+^ T cell and B cell populations in the prostates of h*SKP2*-KI mice. Meanwhile, SKP2 overexpressing prostate organoids were developed for convenient and efficient screening of selective SKP2 inhibitors, which is validated by testing Flavokawain A (FKA), a naturally occurring chalcone in the Kava plant that acts as a SKP2 degrader, as we previously reported [17]. Consistent with our previous findings, FKA significantly inhibits growth of SKP2 overexpressing prostatic organoids, while having minimum toxicity on the normal prostate organoids. Our results suggest that the humanized h*SKP2*-KI mouse model and the organoids derived from the model both are valuable tools to study prostate cancer.

## Materials and Methods

### Establishment of prostate-specific human *SKP2* knock-in mouse model

P2A-Human *SKP2* coding sequence was knocked into the endogenous mouse *probasin* at *exon 1* locus by a CRISPR/Cas9-based approach by the University of California Irvine Transgenic Mouse Facility. After sequencing, heterozygous h*SKP2*-KI females were crossed with wild-type FVB/N male mice to generate male offspring specifically expressing h*SKP2* in all lobes of mouse prostates. Mice were identified with ear tagging and the tail DNA was subjected to polymerase chain reaction (PCR) with the following primers: primer 845 (5’-ATACCTGAAACATGGGATAGGCAC-3’), 846 (5’-TTGCTACTCAGGTCTGGAATCTC-3’) and 847 (5’-ACCAAGTTCCAGAACATTCGTTTC-3’). Prostate lobes of h*SKP2*-KI and wild type (WT) male mice at different ages (4, 9 and 14 months) were dissected, weighed and fixed with 10% neutral formalin for histological and immunohistochemistry analyses or snap frozen for RNA and protein extraction. Prostate-specific expression of human *SKP2* was confirmed by quantitative PCR, Western blotting, and immunohistochemical staining. Histological evaluation of mouse prostatic lesions was performed according to the publication by Dr. Cardiff [18]. Animal care and experiments were carried out according to institutional guidelines and the approved protocol by University of California, Irvine (protocol #: AUP□19□109).

### Quantitative PCR assay

Total RNAs were extracted immediately from the dissected prostate lobes of WT and h*SKP2*-KI mice and human prostate cancer cell lines of PC3/cDNA3.1 and PC3/SKP2 using RNAStat (Amsbio, Friendswood, TX). The primer sequences were designed using PrimerBank (https://pga.mgh.harvard.edu/primerbank/) and the specificity of the primers was checked using nucleotide BLAST in the NCBI. The primer sequences were listed in supplementary table 1.

Primers used for the quantitative PCR were purchased from OriGene (Rockville, MD) or synthesized by Eurofins Genomics (Louisville, KY). Oligo-dT GoScript^TM^ Reverse Transcription mix (Promega, Mandison, WI) was used for the synthesis of the first-strand cDNA. The iTaq SYBR green super-mix (BioRad, Hercules, CA) was used in the real-time PCR reaction on CFX Connect Real-Time System (BioRad, Hercules, CA). Relative quantitative fold changes compared to control were calculated using the comparative 2^−ΔΔCt^ method [19]. *Gapdh* and *ACTB (*β-ACTIN) gene were used as internal controls.

### Western blot analysis

Prostate tissues from the prostate lobes of WT and h*SKP2*-KI mice were homogenized in radioimmunoprecipitation assay (RIPA) buffer with protease inhibitors, and protein concentrations were determined by Bio-Rad DC protein assay (BioRad, Hercules, CA). Equal amount of total protein was loaded onto denaturing SDS-polyacrylamide gel (8-16%) for electrophoresis and then transferred onto PVDF membranes as described previously [20]. After blocking with 5% non-fat milk, the membranes were probed with antibodies against SKP2 (Abcam Inc, Waltham, MA), p27^Kip1^ (BD Biosciences, Franklin Lakes, NJ), androgen receptor (AR) (Cell Signaling Technology, Beverly, MA), or β-TUBULIN (Santa Cruz Biotechnology, Dallas, TX) overnight at 4°C. The membranes were then incubated with corresponding anti-mouse or anti-rabbit secondary antibodies (Cell Signaling Technology, Beverly, MA) followed by wash and visualization by Prometheus Protein ProSignal® Dura ECL Reagent (Genesee Scientific, El Cajon, CA) on X-ray films.

### Immunohistochemistry analysis

Prostate tissues were fixed in 10% neutral buffered formalin, paraffin embedded and sectioned at 5μm thickness. Sections were deparaffinized, hydrated, followed by antigen retrieval with sodium citrate buffer for 20 minutes in a steamer. The sections were then quenched with 3% hydrogen peroxide for 10 min followed by three washes of PBS and blocked with 3% normal goat serum for one hour. Afterwards, the slides were incubated with primary antibodies against Ki67 (Cell Signaling Technology, Danvers, MA) and SKP2 (Abcam Inc, Waltham, MA), as well as p27^Kip1^ (BD Biosciences, Franklin Lakes, NJ) and with horseradish peroxidase (HRP) labeled secondary antibody. Sections were visualized with 3, 3-diaminobenzidine (DAB) using the Cell and Tissue Staining kit (R&D Systems, Minneapolis, MN) and imaged under Keyence BZ-X710 microscope (Itasca, IL).

The images were quantified in Image J by counting the total number of positively stained cells at 12 arbitrarily selected fields at ×40 magnification in a double-blind manner. Images from 3 mice per group were used for statistical analysis.

### RNA sequencing and analysis

Ventral lobes of prostates from h*SKP2*-KI (n=4) and WT mice (n=4) at 12 months old age were freshly dissected and immediately homogenized in RNAStat (Amsbio, Friendswood, TX). The integrities of total RNA were analyzed by Agilent 2100 with the RNA integrity numbers ranging from 8 to 10. mRNA was purified using poly-T oligo-attached magnetic beads and after fragmentation, the first strand cDNA was synthesized using random hexamer primers followed by the second strand cDNA synthesis. The library was ready after end repair, A-tailing, adapter ligation, size selection, amplification, and purification. Libraries were then checked with Qubit and real-time PCR for quantification and bioanalyzer for size distribution, then they were pooled and sequenced on Illumina Platform PE150 (20 million paired-end reads) by Novogene Corporation Inc (Sacramento, CA). The raw paired-end reads were trimmed and quality checked in CLC Genomic Benchwork (Qiagen, Hilden, Germany), followed by mapping to the mouse genome. Normalized counts were used to analyze the gene expression patterns. The raw and normalized data files are deposited in Gene Expression Omnibus (GEO) with series entry number GSE295398. Differently expressed genes (DEGs) in h*SKP2*-KI mouse prostate samples were identified compared to WT mouse prostate samples with the false discovery rate (FDR) <= 0.05. Gene ontology (GO) and hallmark pathways were analyzed using Gene Set Enrichment Analysis software (GSEA) from the Broad Institute. Enriched pathways were treated as significant with FDR <= 0.05.

### Migration and invasion assay

Falcon® HTS 24-well Multiwell system (Cat# 351185, Corning, AZ) and Corning® BioCoat™ Matrigel® Invasion Chamber (Cat#354480, Corning, AZ) were used for migration assay and invasion assay, respectively, as described previously [21]. After cell starvation for 24 hours, 30,000 PC3/pcDNA3.1 and PC3/SKP2 cells were seeded onto the upper insert in 500µl RPMI1640 medium without serum, and 750 µl RPMI 1640 with 10% FBS was added to the bottom well as chemoattractant or RPMI 1640 without serum as control well. After 72 hours, non-migrating or non-invading cells were removed from the upper surface of the membrane by moistened cotton swab scrubbing followed by cell fixation with cold methanol and staining with hematoxylin. Images from 5 different areas of each insert were acquired and cell numbers of 3 replicate inserts from each cell line were used for statistical analysis.

### Culture of prostate organoids and evaluation of SKP2 inhibitors’ anti-tumor effects

We have followed the protocol that was published by Hans Clevers’ group [22]. Fresh prostate tissues from *SKP2*-KI and WT male mice were minced into small pieces and digested in 5 mg/ml Collagenase II (Life Technologies, Carlsbad, CA) solution containing 10μM Y-27632 (AbMole BioScience, Houston, TX) for 2 hours at 37°C with shaking. After washing with adDMEM/F12+/+/+ medium containing penicillin/streptomycin, 10mM HEPES, and 1X GlutaMAX (Life Technologies, Carlsbad, CA), the pellets were further digested in TrypLE solution (Life Technologies, Carlsbad, CA) containing 10μM Y-27632 for approximately 15 minutes at 37°C with pipetting every 5 minutes, followed by washing and centrifugation. 20,000 digested cells were resuspended in cold Matrigel and plated 20μl/well into prewarmed 48 well plates. The plates were then upside down in the 37°C incubator for 15 min to allow the Matrigel to solidify. 300μl-500μl pre-warmed mouse prostate culture medium plus 10μM Y-27632 was added into each well. Culture medium was refreshed every 2-3 days. TrypLE solution supplemented with Y27632 was used for passaging the organoid culture.

WT and h*SKP2*-KI prostate organoids were fixed with 4% paraformaldehyde, embedded in HistoGel (Thermo Scientific, MA) and sectioned for H&E staining for histological evaluation and for immunohistochemistry analysis of AR, SKP2 and p27^Kip1^ expression.

To evaluate the effects of known SKP2 inhibitors FKA and C1 on the growth of prostate organoids, hSKP*2*-KI prostate organoids and WT prostate organoids were plated into 96 well plates and then treated with vehicle control (0.5% DMSO) or FKA or C1 inhibitor in triplicate. The morphology of the organoids was imaged before the treatments and at different time-points after the treatments under Keyence fluorescence microscopy. The size of each organoid was analyzed by Image J software. In addition, the viability of the organoids was determined by measuring fluorescence intensities from the conversion of a cell-permeant nonfluorescent Calcein AM dye (CalAM, Thermo Scientific, MA) into a green fluorescent Calcein (ex/em 495 nm/515 nm) in living cells. The fluorescence images were obtained at different treatment times under Keyence fluorescence microscopy and analyzed by Image J. Fluorescence intensity and size of each organoid acquired at different time points were compared to its intensity and size at day 0 to obtain the relative changes over time (n=15 for WT or KI prostate organoids).

#### Cell deconvolution of bulk RNA-seq data

Mouse prostate single cell signature was obtained from the study by Joseph DB et al [23]. The mouse prostate single cell annotation was downloaded from GUDMAP database at https://doi.org/10.25548/17-DRBC, The rds file was processed in R software and mouse prostate single cell genes and cells annotation matrix were extracted as the signature file used in CIBERSORTx (https://cibersortx.stanford.edu/). Bulk RNA-seq data from WT and h*SKP2*-KI mouse prostates were used as a mixture input according to the CIBERSORTx tutorial. S-mode batch correction was applied and estimation of cell fractions from bulk tissue transcriptomes was generated on CIBERSORTx as described in published paper [24].

### Statistical analysis

Student’s t-test, one-Way ANOVA followed by the Bonferroni t-test for multiple comparisons and log-rank testing for survival analysis were carried out using GraphPad Prism 9.0 to calculate statistical significance. Data are expressed as mean ± standard deviation (SD). All statistical measures were two-sided, and *p*-values <0.05 were statistically significant.

## Results

### Establishment of prostate specific human *SKP2-*KI mouse lines by a CRISPR/Cas9 method

The P2A-h*SKP2* coding sequence was introduced into the guide RNA generated DNA break by a CRISPR/Cas9 method in the endogenous mouse *probasin* at exon 1 locus (Fig. 1A) for expression of h*SKP2* under control of the mouse *probasin* promoter. Probasin-h*SKP2*-KI mouse lines were screened using specific PCR of genomic DNA extracted from mouse tail biopsy and confirmed the correct sequencing of the genomic insert by the Sanger sequencing. Nineteen of 43 (44.2%) KI founder mice were genotyped positive. Three probasin-h*SKP2*-KI mouse lines (L9929, L10055 and L10092) show 5, 7-to-35-fold overexpression of h*SKP2* in the prostate compared to that in the prostate of wild-type mice (Fig.1B). Western blotting analysis using human specific anti-SKP2 antibody (Abcam #ab68455) reveals that human SKP2 protein is expressed in the microdissected prostate lobes (anterior, dorsal, lateral, and ventral lobes) of three probasin-hSKP2-KI mouse lines but not in the prostate of wild-type mice (Fig. 1C). The expression level of hSKP2 protein is lower in the anterior lobe compared to other prostate lobes (Fig. 1C). Consistently, p27^Kip1^ protein, a putative substrate of SKP2 is down-regulated in the prostate of probasin-h*SKP2*-KI line L10092 (Fig. 1D). There is no detectable hSKP2 protein in other organs of the L10092 line, including testis, kidney, spleen, and liver (Fig. 1E). These results indicate that SKP2 overexpression is prostate-specific in the probasin-h*SKP2*-KI lines.

**Figure 1.**
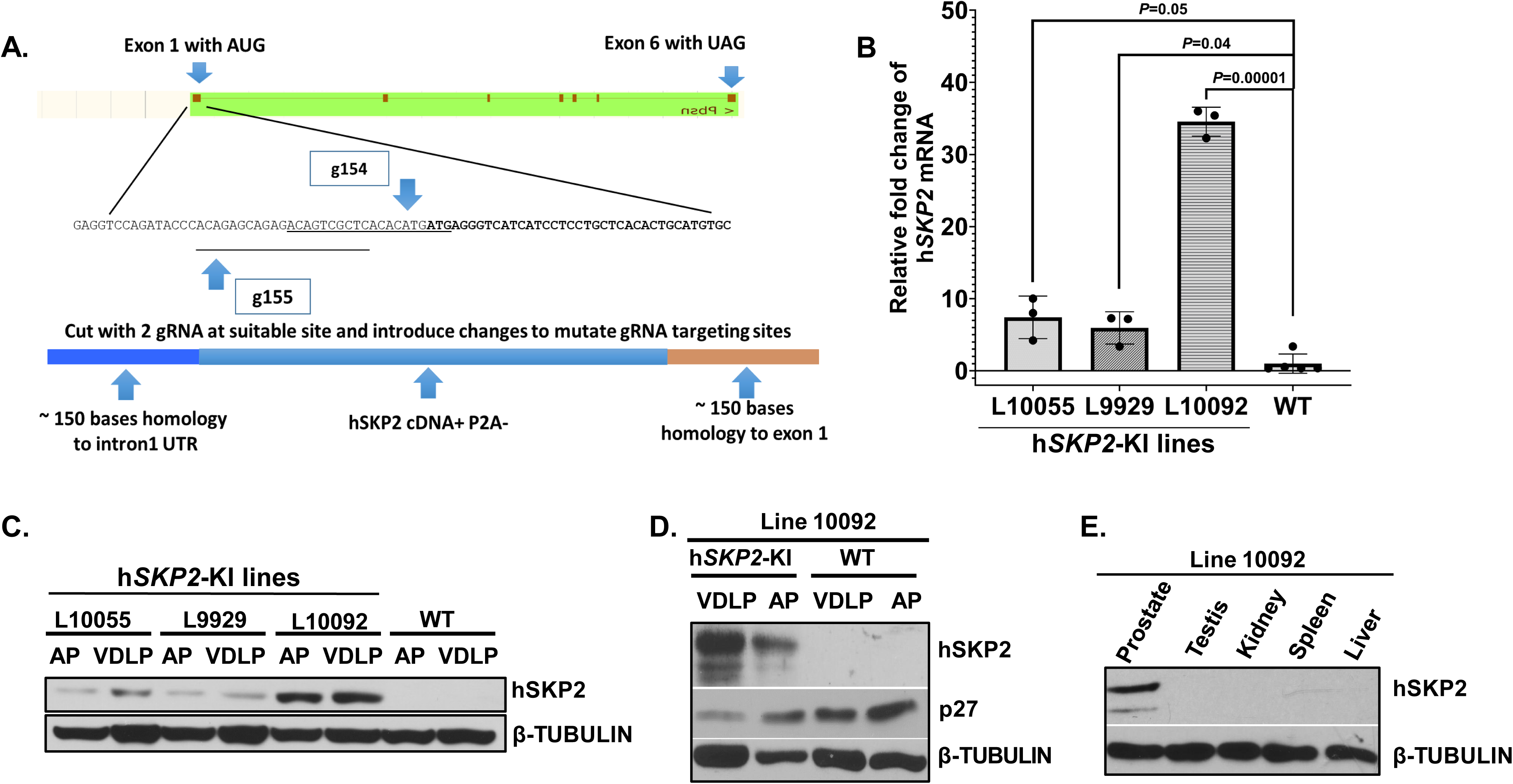
Establishment and characterization of prostate-specific SKP2 humanized mice. **A.** The Scheme of h*Skp2* KI allele. The P2A-h*SKP2* sequence was introduced into the gRNA generated DNA break in mouse *probasin* locus, which is specifically expressed in the prostates of a mouse. **B.** Mice were genotyped, sequenced and three founder lines (L10055, L10092, and L9929) of h*SKP2* KI mice were obtained. The mRNA expression levels of h*SKP2* in the prostate of WT (wild-type) *v.s.* h*SKP2*-KI mice were determined by the real-time PCR method. N=5 for WT mice and n=3 for each KI line. **C.** SKP2 protein expression levels in ventral, dorsal, and lateral (VDLP) and anterior (AP) prostate lobes of WT and h*SKP2* KI mice were examined by Western blotting analysis. **D**. The protein expression of p27^Kip1^ in prostate lobes of WT and h*SKP2*-KI mice. **E.** Expression levels of hSKP2 protein were compared among the prostate, testis, kidney, spleen and liver of h*SKP2* KI mice. Mice were 3-4 months old with n=3 for each tissue examined.

### SKP2 overexpression induces hyperplasia, mPIN and low-grade carcinoma associated with increased SKP2 expression and cell proliferation and decreased expression of p27^Kip1^

Histological evaluation shows that H&E-stained prostate lobes (anterior, dorsal, lateral, and ventral lobes) of probasin-h*SKP2*-KI mice at different ages (4, 9, and 14 months of age) exhibit different grades and proportions of marked enlargement and variation in the duct size, thicker and disoriented epithelial layers, primitive cribriform patterns, filled duct lumens, vacuolated cytoplasm, prominent round to oval nuclei with one or more nucleoli, and multiple mitotic figures, while similar lesions were not found in the prostate of wild type mice at the same ages (Figs. 2A & B). At 4 months of age, 63.3 % (7 out of 11) of probasin-h*SKP2*-KI mice were observed with hyperplasia and low-grade mPIN; 50% (6/12), 42% (5/12), and 8% (1/12) of probasin-h*SKP2*-KI mice with low-grade PIN, high-grade PIN, and hyperplasia, respectively, at 9 months of age; 6.7% (1/15), 80% (12/15) and 13.3% (2/15) of probasin-h*SKP2*-KI mice with low grade PIN, high grade PIN and low-grade carcinoma at 14 months of age, respectively (Fig. 2C). The mean prostate weights of probasin-h*SKP2*-KI mice were significantly increased compared to those of wild-type mice (P < 0.05) at 14 months of age (Fig. 2D).

**Figure 2.**
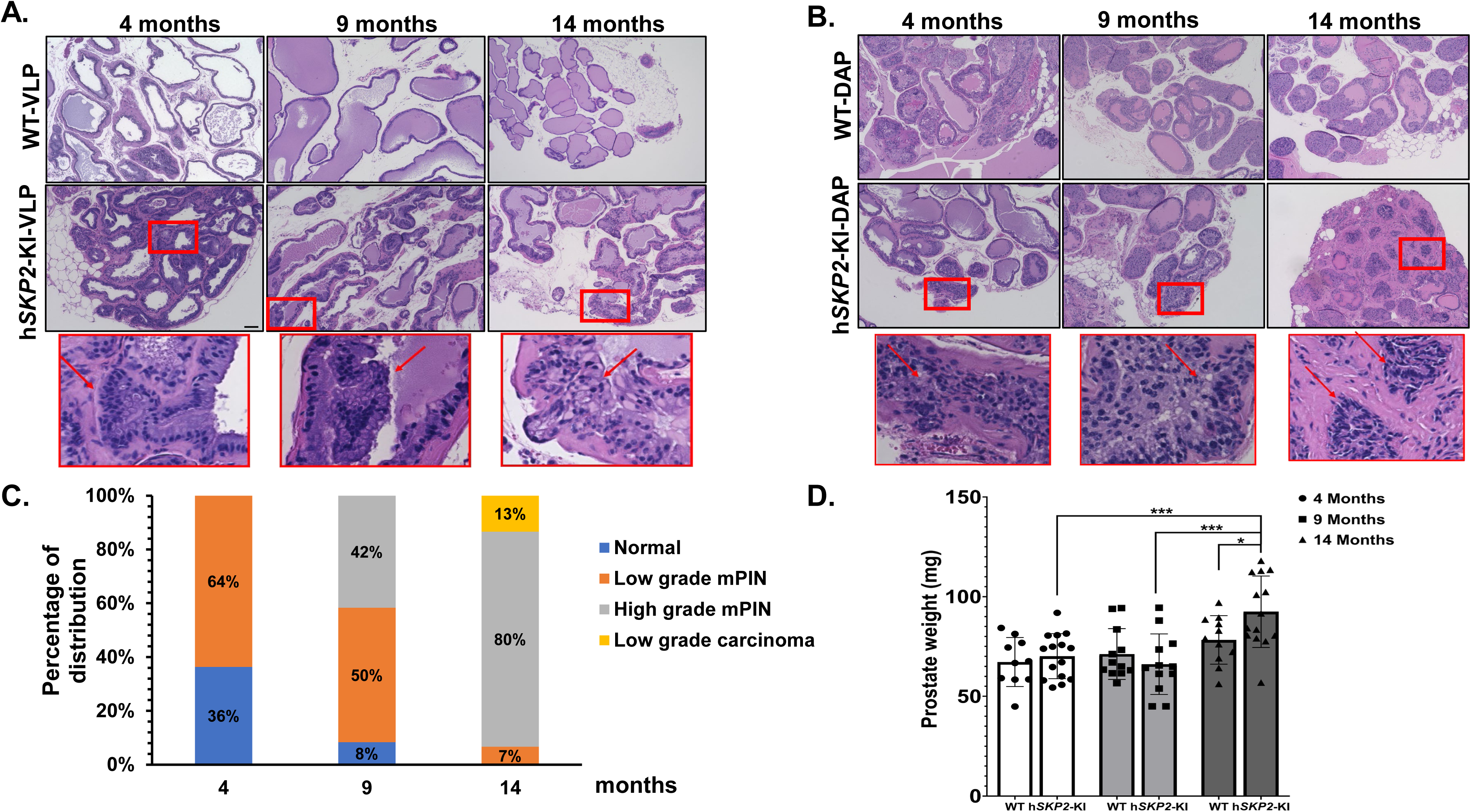
Histopathological evaluation of the prostate of h*SKP2*-KI and WT mice. **A. & B.** Representative images of H&E stained sections from ventral, dorsal, and anterior prostate glands of WT and h*KP2*-KI mice are shown. Overexpression of human SKP2 in the prostate of mice results in hyperplasia at 4 months of age. Increased layers of epithelial cells were observed (arrow) in prostate lobes of hSKP2-KI mice, whereas the prostate epithelium of WT mice is in flat layer and maintains normal columnar and cuboidal shape. Mouse prostatic intraepithelial neoplasia lesions were observed in h*SKP2*-KI mice at age of 4, 9 and 14 months, featured by nuclear stratification, enlargement and hyperchromasia. Low-grade carcinoma was detected in the prostate of hSKP2 mice at 14 months of age. Atypical epithelial cells proliferate and partially fill the duct. Enlarged nuclei and prominent macronucleoli present in the nests of cells indicated low grade carcinoma. Scale bar: 100 µm. **C.** The distribution of prostatic lesions in h*SKP2*-KI mice at age of 4 (n = 11), 9 (n = 12) and 14 (n = 15) months. **D.** Prostate weights of h*SKP2*-KI and WT mice were measured at 4 (n = 15 and 10), 9 (n = 12 and 12), and 14 (n = 14 and 11) months of age, respectively. A significant increase in the mean prostate weights was observed in h*SKP2*-KI mice at 14 months of age compared to that of WT mice (\**P* < 0.05, \*\*\**P* <0.001).

Immunohistochemistry analysis reveals that strong SKP2 positive staining was mainly observed at the sites of prostatic lesions (i.e. hyperplasia, PINs and low-grade carcinoma), accompanied by decreased staining intensities of p27^Kip1^ in probasin-h*SKP2*-KI mice compared to those from wild-type mice (Fig. 3A). In addition, cell proliferation was significantly enhanced in the prostates of probasin-h*SKP2*-KI mice as indicated by increased Ki67 positive staining cells by approximately 26 folds compared to those from wild-type mice (P < 0.01) (Figs 3B & D), while there is no significant difference in AR expression in the prostates between probasin-h*SKP2*-KI mice and wild-type mice (Figs. 3C & E).

**Figure 3.**
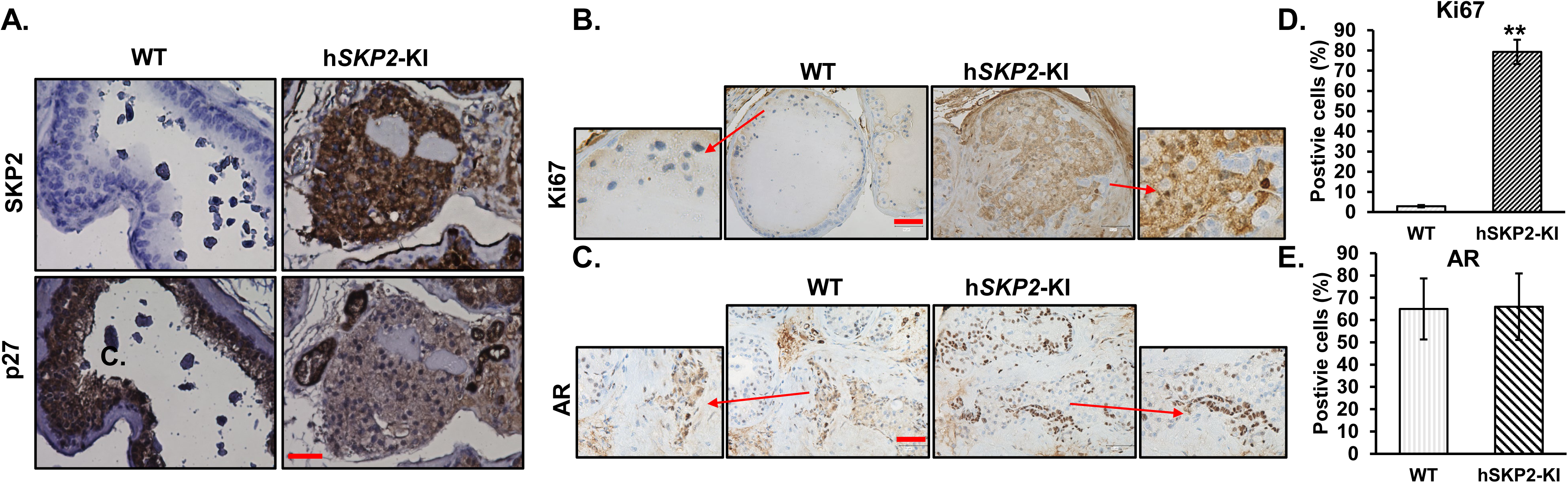
Examination of hSKP2, p27^Kip1^, and AR expression by immunohistochemistry analysis of WT and h*SKP2*-KI mouse prostates. **A., B., and C.** Immunohistochemistry staining reveals robust expression of human SKP2 protein in the prostate of h*SKP2*-KI mice accompanied by a reduction in p27^Kip1^ expression. Representative images of hSKP2. P27^Kip1^, Ki67 and AR staining on the prostate of WT and h*SKP2*-KI mice. Scale bar: 20 µm for hSKP2 and p27Kip1 and 50 µm for Ki67 and AR staining images. **D. & E.** Ki67 and AR positively stained cells were counted in 12 arbitrary fields under microscope. The mean percentage of Ki67 positive cells was significantly elevated in h*SKP2*-KI mouse prostates compared to WT mouse prostates. The mean percentages of AR positive cells are not significantly different between WT and h*SKP2*-KI mice prostates. \*\**P*<0.01. Mice used for the immunohistochemistry study were 12-14 months old and n=4.

### Gene expression profiling reveals differentially expressed genes and significant enrichment of EMT and interferon pathways in SKP2 overexpressing mouse prostates

We performed bulk RNA sequencing to determine the effect of SKP2 overexpression on global gene expression alterations in the prostate of probasin-h*SKP2*-KI mice and wild-type mice. A total of 1,753 differentially expressed genes (DEGs) were identified with a false discovery rate (FDR) *p*-value < 0.05. Among them, 678 genes demonstrated more than a two-fold change in expression, including 498 genes being up-regulated and 180 genes downregulated (Figs. 4A & B). The top DEGs include *Fcgbp, Krt4, Slc6a2, Mme, Egf, Adh6a, Clu, Lrrc31, Sim2, Mmp12, Ly6k, Ly6d, Ly6a, etc*. Androgen responsive genes (*Ccnd1, Hmgcr, Tmprss2, Insig1, Dhcr24*, and *Bmpr1b),* G2M related genes (*Slc7a1, Egf*, and *Cul4a*) and apoptosis related genes (*Clu*, *Bcl2*, and *Sc5d*) are also on the top list of DEGs. The Gene Ontology (GO) gene set enrichment analysis (GSEA) highlights the activation of extracellular matrix components, leukocytes migration, and mesenchymal cell proliferation pathways in SKP2 overexpressing mouse prostates (Fig. 4C). In addition, GSEA of hallmark genes revealed that EMT, interferon alpha and gamma, angiogenesis, and inflammatory response pathways pathway are significantly enriched in SKP2 overexpressing mouse prostates (Fig. 4D).

**Figure 4.**
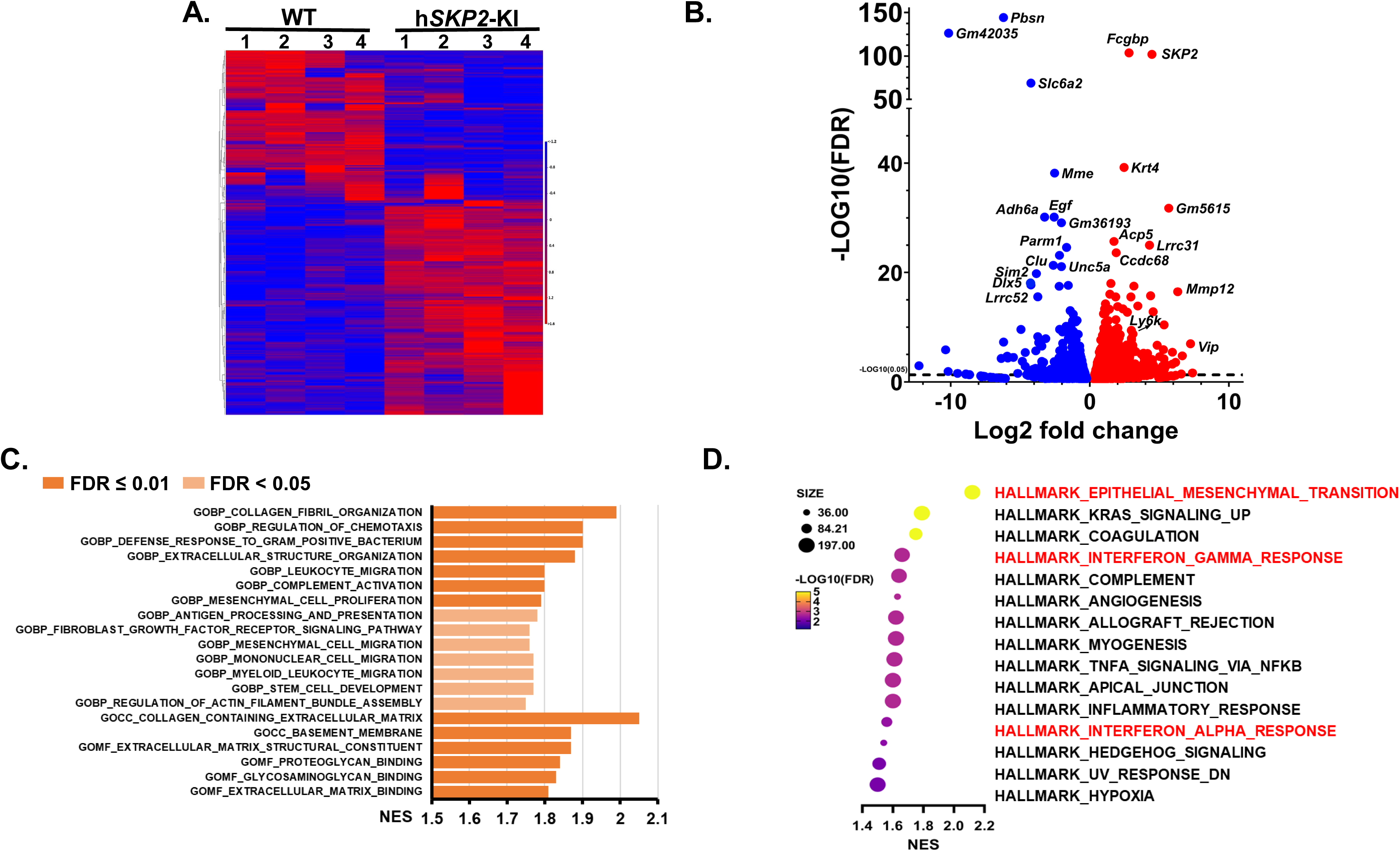
DEGs and significantly altered pathways in the prostate of h*SKP2*-KI mice were unveiled by bulk RNA sequencing analysis. **A.** Hierarchical clustering heatmap of differential gene expressions between WT and h*SKP2*-KI mouse ventral prostates (n=4). **B.** Volcano plot was used to visualize the distribution of significantly upregulated and downregulated genes in h*SKP2*-KI versus WT mouse prostates with an FDR <0.05. **C.** Significantly upregulated components and pathways in h*SKP2*-KI mouse prostates compared to WT mouse prostates by GSEA enrichment of Gene Ontology (GO). **D.** GSEA of hallmark pathways identified marked elevations in pathways associated with EMT and interferon responses in h*SKP2*-KI mouse prostates.

The list of enriched gene expression alterations in EMT and interferon pathways is shown as enrichment profiles and heatmaps in Figs. 5A, B, D and E. The increased gene expression levels of selected EMT pathway genes, including *Fmod*, *Thy1*, *Wnt5a,* and *Vim*, and the down-regulated interferon gamma pathway genes, such as *Tnfaip2*, *Cd38*, *Pfkp*, *Il15*, *Lats2* were verified by quantitative PCR method (Figs. 5C & F).

**Figure 5.**
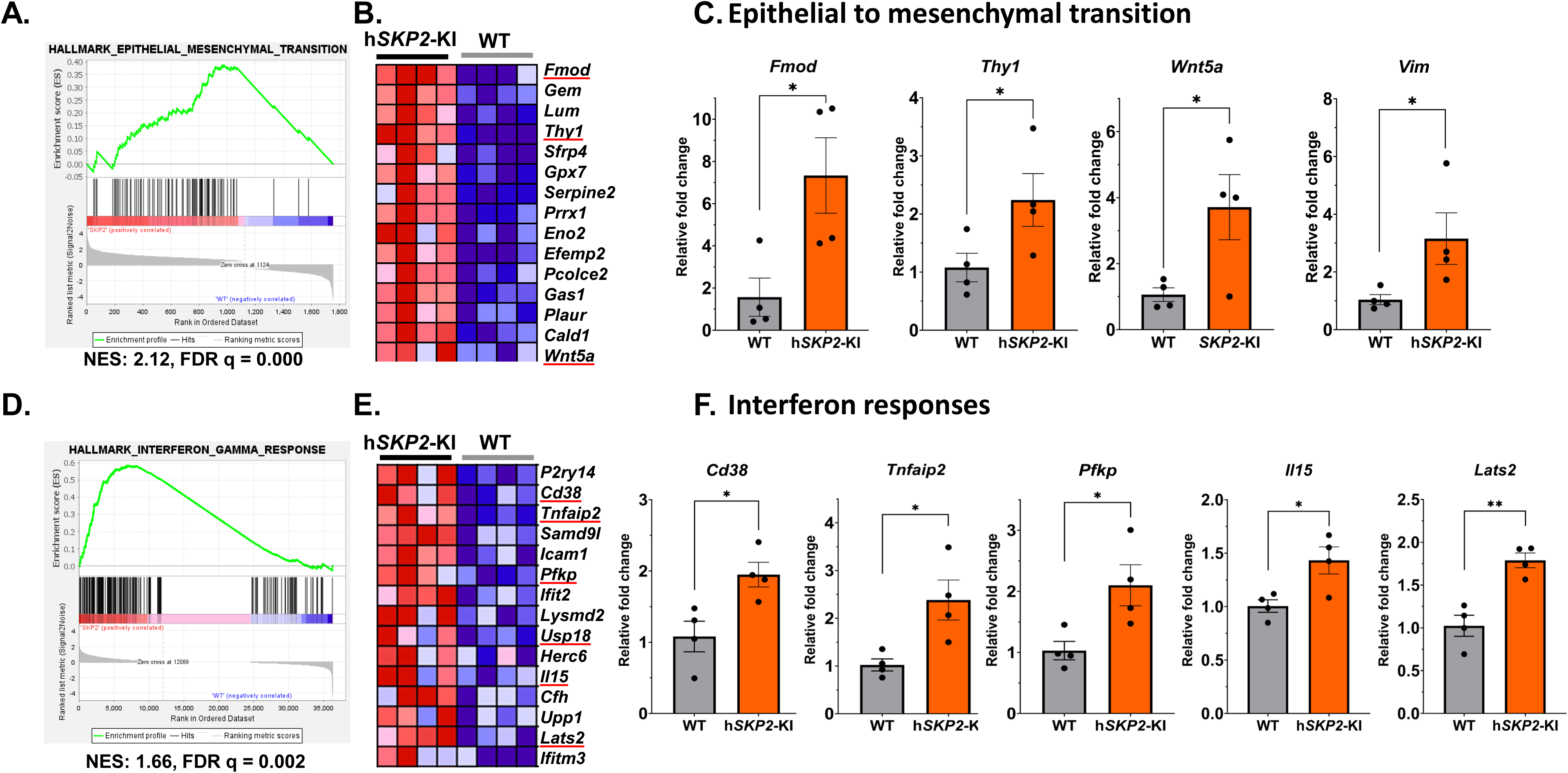
The alterations of EMT and interferon response pathways were highly enriched in h*SKP2*-KI compared to WT mouse prostates. **A.** EMT GSEA enrichment plot. **B.** The heatmap of top enriched genes in the EMT pathway. **C**. Quantitative PCR was performed to verify the expression of EMT-associated genes in mouse prostates, including *Fmod, Thy1, Wnt5a, and Vim*. **D.** GSEA enrichment plot interferon gamma response. **E.** The heatmap of top enriched genes in the interferon gamma response pathway. **F.** Quantitative PCR was used to validate the expression of interferon-responsive genes in mouse prostates, including *Cd38, Tnfaip2, pfkp, Il15,* and *Lats2*. N=4, \**P*<0.05, \*\**P*<0.01.

### Cell deconvolution analysis reveals an increase in fibroblasts and a decrease in infiltrating T and B cells in SKP2-overexpressing mouse prostates

Given the profound alterations in gene expression of EMT, extracellular matrix and interferon pathways in SKP2 overexpressing mouse prostates, we have deconvoluted the probasin-h*SKP2*-KI and WT bulk RNA-seq data using the R package CIBERSORTx. The cell deconvolution analysis revealed significant differences in the proportional compositions of various immune and stromal cell types in the prostate of the h*SKP2*-KI versus WT mice (Fig. 6A). The h*SKP2*-KI mice exhibit a significantly higher proportion of fibroblasts in the prostate compared to that of WT mice (P = 0.0122, Student’s t test) (Fig. 6B). A reduction in lymphoid lineage cells and an expansion of myeloid lineage cells have also been observed in the prostate of h*SKP2*-KI mice compared to that of WT mice. Notably, h*SKP2*-KI mouse prostate glands demonstrate a significantly lower proportion of CD8^+^ T cells, B cells, and myeloid dendritic type I cells compared to WT mouse prostate glands (Ps = 0.0435 to 0.0012, Student’s t test). Proliferating classical monocytes that have the potential to differentiate to tumor-associated macrophages, which suppress T cell function, [25] were found to infiltrate into the prostate of h*SKP2*-KI mice more than that of WT mice (P = 0.008, Student’s t test).

**Figure 6.**
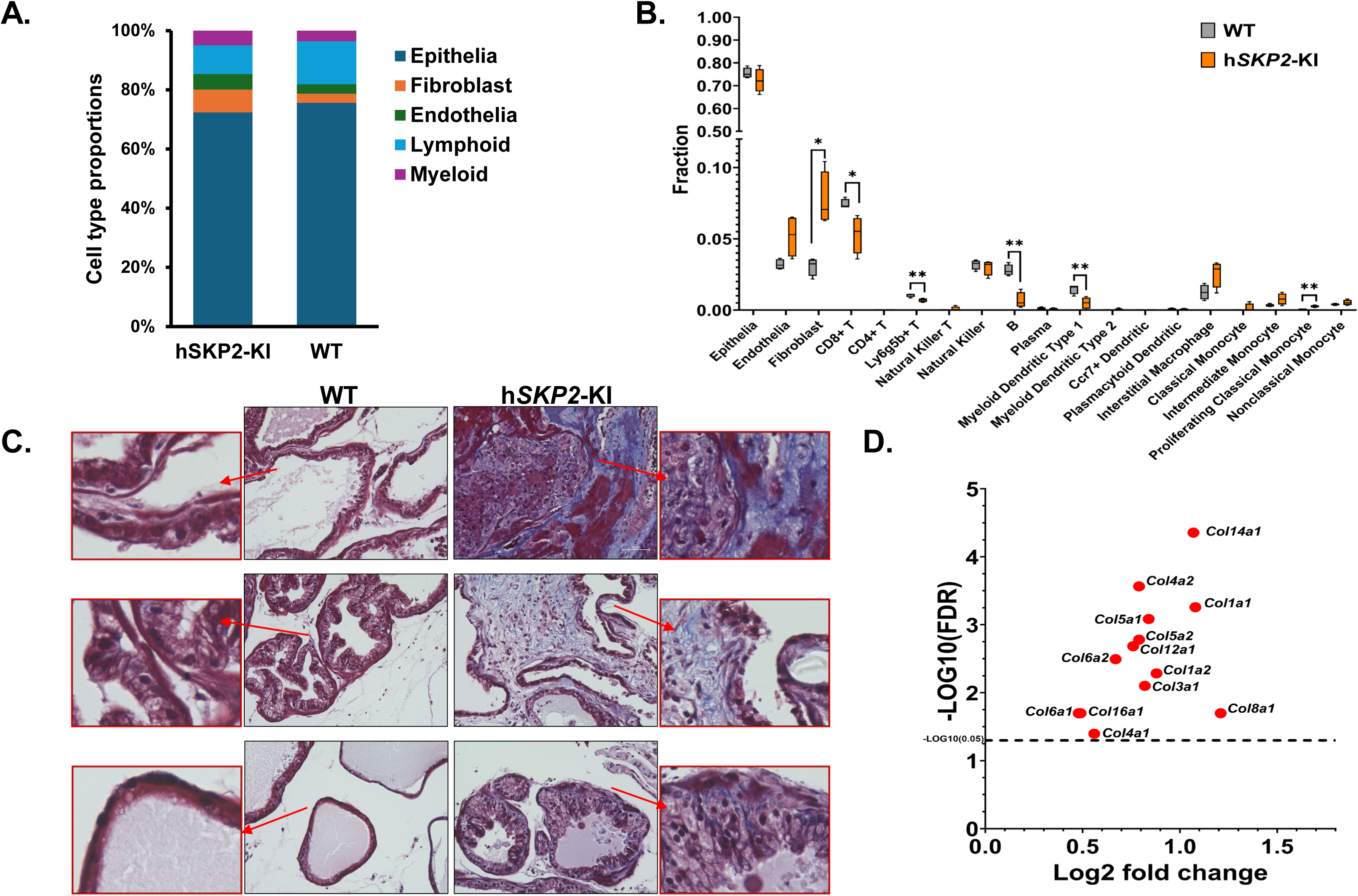
Single cell deconvolution of bulk RNA sequencing data and examination of extracellular matrix by Masson’s trichrome staining analysis in WT and h*SKP2*-KI mouse prostates. **A. & B.** Cell lineage proportions and fractions of different cell types were derived by deconvoluting bulk RNA sequencing data that were generated from WT and h*SKP2*-KI mice prostates. **C.** Masson’s trichrome staining highlights significantly increased collagen deposition in the prostate lobes of h*SKP2*-KI mice compared to those of WT mice, as indicated by stronger blue staining. Scale bar: 50 µm**. D.** Expression of many collagen-related genes is significantly upregulated in the prostates of h*SKP2*-KI mice compared to those of WT mice.

We have further evaluated the extracellular (*i.e.* collagen) components of h*SKP2*-KI and WT mouse prostates using Masson’s Trichrome staining and expression of collagen related genes. Figures 6C and 6D show a markedly enhanced collagen staining and upregulation of numerous collagen related genes in the prostate of h*SKP2*-KI mice versus WT mice. Taken together, these results suggest that SKP2 overexpress induces the remodeling of ECM and creates an immune suppressive prostatic microenvironment to support tumor initiation and development.

### SKP2 overexpression is associated with aggressiveness of human prostate cancer, promotes cell migration and invasion, and regulates the expression of EMT, and genes related to interferon signaling

Next, we explored the cBioportal datasets for the frequencies of *SKP2* genetic alterations (i.e. amplification, deep deletion, and mutation) in cancers [26–27]. We observed that amplification of *SKP2* gene frequently occurs in various cancers, including sarcoma, urothelial or bladder cancer, lung cancer, gastric cancer, metastatic prostate cancer, and prostate adenocarcinoma with a range from about 12% to 6% (Fig. 7A). Analysis of RNA expression levels in prostate tumor tissues in the TCGA dataset also demonstrates that higher levels of *SKP2* mRNA expression (>= 2 fold) are more frequently associated with aggressive prostate tumors with higher Gleason scores compared to lower Gleason score prostate tumors (11% and 75%for Gleason score 9 and 10, respectively versus 3 to 1% for Gleason score 8 to 6) (Fig. 7B). Further analysis of RNA expression levels with available survival data in the SU2C/PCF Dream Team, PNAS 2019 study shows that higher levels of mRNA levels in prostate tumor tissues are significantly associated with poorer survival of prostate cancer patients (P < 0.05, Log-rank test). These results indicate that SKP2 plays a significant role in the development of advanced prostate cancer and suggest that *SKP2* overexpression is an unfavorable prognostic factor associated with disease aggressiveness.

**Figure 7.**
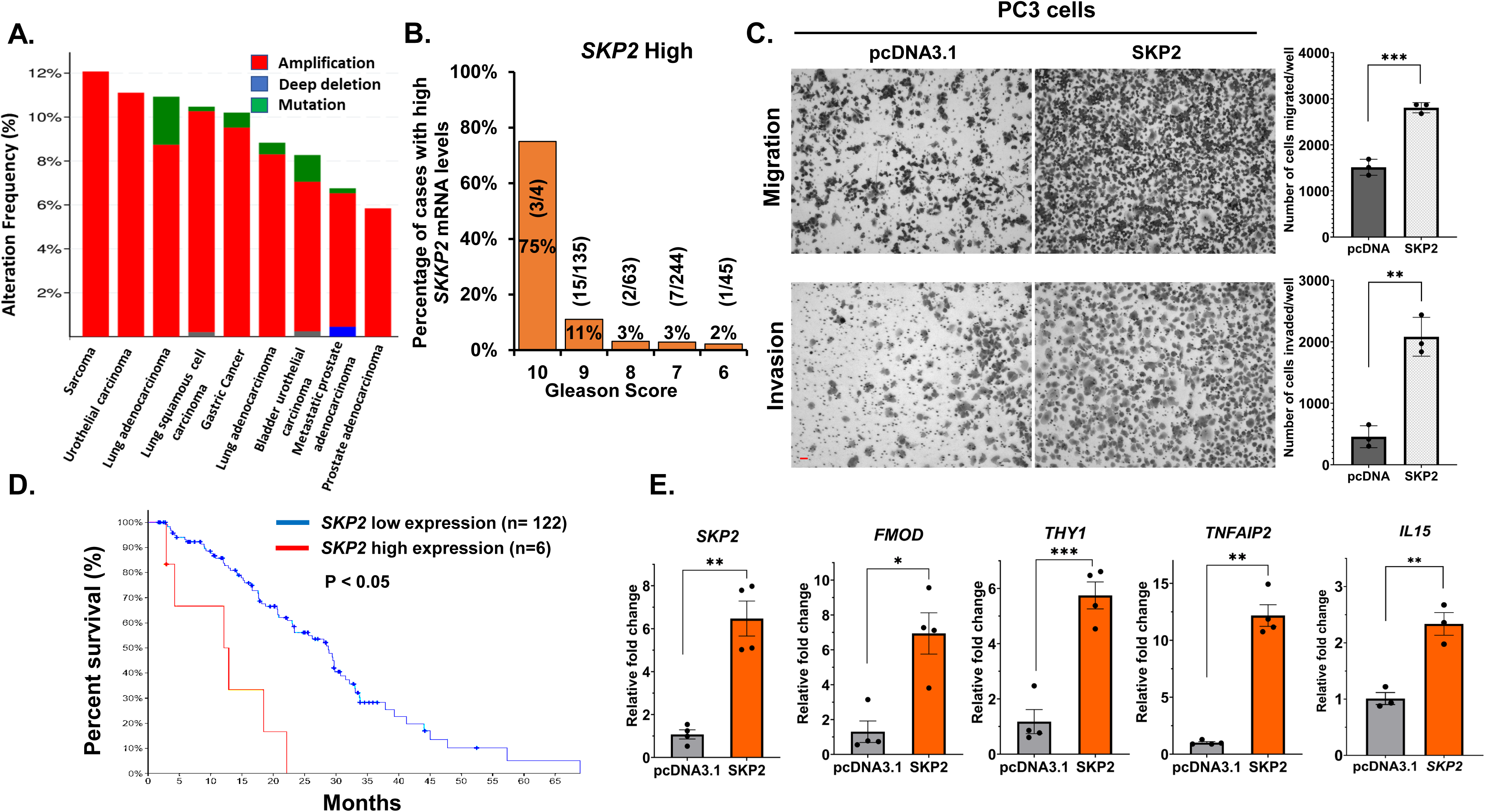
SKP2 Overexpression in human prostate cancer is associated with the aggressiveness of prostate cancer through promoting cell migration and invasion, and regulating EMT and interferon signaling. **A.** human *SKP2* gene is frequently amplified in multiple cancers in cBioportal datasets. **B.** Prostate cancer patients with a Gleason score of 9 or 10 are more likely to have a higher SKP2 mRNA level in their tumor tissues (data were derived from the TCGA dataset). **C.** Prostate cancer PC3 cells overexpressing SKP2 exhibit significantly enhanced migratory and invasive capabilities compared to PC3 cells expressing vector control. Scale bar: 50 µm**. D.** Higher SKP2 mRNA expression levels in prostate tumor tissues are significantly associated with poorer survival of prostate cancer patients with metastatic disease (Log-rank test, P < 0.05, data were from SU2C/PCF Dream Team, PNAS 2019). **E.** The overexpression of SKP2 in PC3 cells led to the upregulation of genes involved in EMT and interferon responses related genes. N=4, statistical significance is indicated as \**P*<0.05, \*\**P* <0.01 and *** *P* <0.001.

To further evaluate the association of SKP2 overexpression with aggressiveness of prostate cancer, we have performed cell migration and invasion experiments, as well as quantitative PCR analysis of the expression of EMT and interferon pathways related genes using a prostate cancer cell line PC3 with or without SKP2 overexpression. Figure 7 shows that SKP2 overexpression increases cell migration and invasion of PC3 cells by 1.8 and 4.5 folds, respectively, compared to PC3 cells expression vector control pcDNA3.1 (Ps < 0.01, Student’s t test). In addition, SKP2 overexpression in PC3 cells increases the expression of EMT and interferon gamma pathway related genes as described above in the h*SKP2*-KI mouse model, including *FMOD*, *THY1*, *TNFAIP*, *IL15*, *etc.* (Fig. 7E and supplementary Fig.1). Taken together, data from the h*SKP2*-KI mouse model, human prostate cancer clinical data and in vitro human prostate cancer cell culture studies are consistent, at least in part, to support the critical role of SKP2 overexpression promote aggressiveness of prostate cancer by regulating the EMT and interferon pathways. Our data suggest that the humanized mouse model recapitulates human prostate cancer.

### Generation and characterization of prostate organoids derived from h*SKP2*-KI mice for testing SKP2-targeting agents

Since in vivo studies of SKP2 targeting agents using mouse models are often time consuming and cost-prohibitive, we therefore established cultures of prostate organoids from the prostate of both h*SKP2*-KI and WT mice to facilitate screening of selective SKP2 targeting agents. H & E staining and histological analysis shows that the morphology of prostate organoids from WT mice is characterized by a branched, acinar-like structure and mimics the in vivo epithelial architecture of mouse prostate tissue, while prostate organoids from h*SKP2*-KI mice display a morphology of a distorted acinar-like structure with multiple layers of cells and filled lumens (Fig. 8A). Prostate organoids from h*SKP2*-KI mice also show strong expression of SKP2 accompanied by negative staining of p27^Kip1^ whereas prostate organoids from WT mice exhibit the opposite expression status of SKP2 and p27^Kip1^. AR expression is also positive in prostate organoids derived from both *SKP2*-KI and WT mice (Fig. 8A).

**Figure 8.**
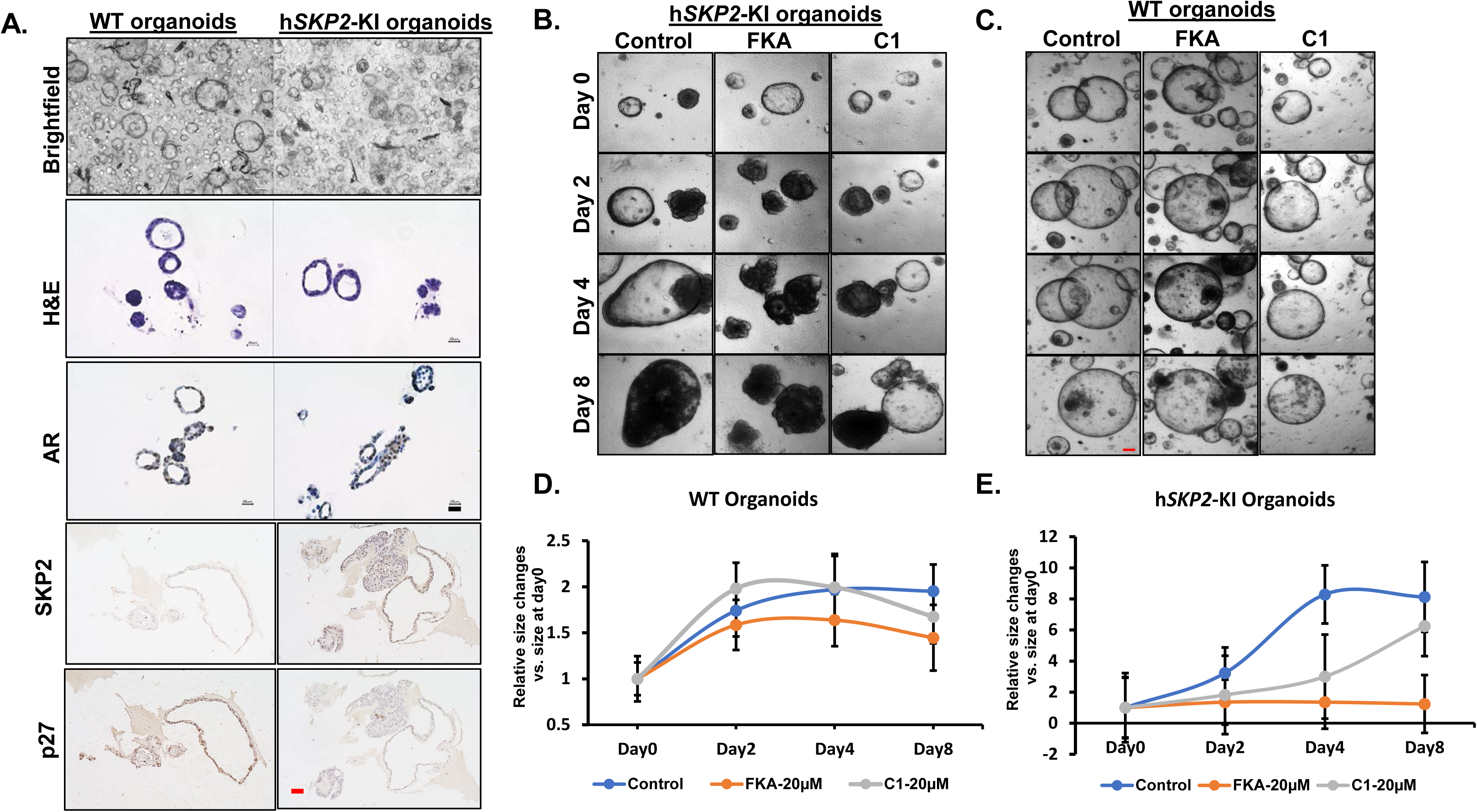
Characterization and validation of mouse prostate organoids derived from WT and h*SKP2*-KI mouse prostates as a tool to test SKP2 targeting agents. **A.** Mouse prostate lobes were dissected, digested into single cells, and seeded in Matrigel. Organoids formation was visualized by a brightfield microscope after seeding for 5 to 7 days. Prostatic organoids derived from WT and h*SKP2*-KI mice recapitulate prostate acinar architecture and express AR, as shown by H&E and immunohistochemistry staining, respectively. Scale bar: 20 µm. Human *SKP2*-KI organoids exhibit stronger SKP2 staining but weak p27Kip2 expression compared to WT organoids. Scale bar: 50 µm**. B. & C.** Representative images of WT and h*SKP2*-KI mouse prostate organoids that were treated with FKA and C1 at 20µM for 8 days. **D. & E.** These organoids were imaged and recorded every two days and the size of these organoids were calculated by image J software. Scale bar: 100 µm.

We have previously reported that FKA acts as a SKP2 degrader for selectively inhibiting the growth of prostate cancer PC3 cells overexpressing SKP2 [17]. In addition, SKP2 inhibitor C1 has been shown to specifically target the interaction between SKP2 and p27^Kip1^, preventing the binding and subsequent ubiquitination and degradation of p27^Kip1^ [28]. To validate our organoid model, we therefore tested the selectivity of these known SKP2 inhibitors on the growth of SKP2 overexpressing prostate organoids versus normal prostate organoids to establish a “proof of concept” tool for future screening with more SKP2 inhibitors. Figures 8 B & C show that control-treated h*SKP2*-KI organoids continuously increase their sizes until day 4 after seeding, whereas FKA- and C1-treated h*SKP2*-KI organoids exhibit a significantly slower growth in size (Ps <0.05). However, C1-treated h*SKP2*-KI organoids recovered from its growth inhibition on day 8, whereas the size of FKA treated h*SKP2*-KI organoids continued to decrease. This result suggests that h*SKP2*-KI organoids could potentially develop resistance to C1. In addition, both C1 and FKA exhibit no significant growth inhibitory effects on wild-type prostate organoids.

FKA at 12.5 μM completely reduced cell viabilities of hSKP*2*-KI organoids 3 days after the treatment but had no effect on the growth of wild-type organoids (Supplementary Fig. 2). FKA exhibited excellent selectivity to the growth of hSKP*2*-KI organoids over wild-type organoids.

## Discussion

Our analysis of publicly available datasets shows that *SKP2* gene is frequently amplified in multiple cancers, including sarcoma, urothelial cancer, and prostate cancer, and that higher *SKP2* mRNA levels in prostate tumor tissues are associated with higher Gleason grades and poorer overall survival of prostate cancer patients. In addition, SKP2 interacts with pRB, PTEN, AR, and H-RAS in prostate cancer [4, 15, 29, 30]. Allelic loss or reduced expression of RB occurs in 25% to 50% of human prostate cancer [31–33], and PTEN deletion and/or mutations were detected in 30% of primary prostate cancer and 63% of metastatic prostate cancer [34, 35].

Genetic depletion or pharmacologic inhibition of SKP2 can block the pRB deficient tumorigenesis by inducing apoptosis or suppress cancer progression by triggering p53-independent senescence in prostates of *Arf*^-/-^ mice and PTEN deficient mice [2, 12, 14, 15, 36]. All evidence suggests that SKP2 functions as a key oncogene during prostate carcinogenesis and progression. SKP2 is also associated with the acquired drug resistance like paclitaxel resistance in prostate cancer [37, 38]. The acquired chemoresistance will eventually lead to treatment failure of castration-resistant prostate cancer [36, 37]. Since Skp2-knockout mice are viable and fertile [39], SKP2 may serve as an “Achilles’ heel” type of drug target to effectively prevent or treat pRB, p53 and/or PTEN deficient prostate cancer potentially with less toxicity. Therefore, there is an unmet need to develop a h*SKP2*-KI mouse model for further understanding the oncogenic functions of SKP2 in prostate carcinogenesis and to provide a novel tool for developing novel SKP2 inhibitors.

Previous prostate-specific transgenic mouse studies, such as TRAMP and Hi-Myc models, have used random integration of rat *Pbsn* mini promoter driven oncogenes into mouse genome, which requires considerable time, effort, and funds to screen many independent lines with a lower positive rate. In this study, we have developed the novel CRISPR/Cas9-based approach to knock-in a h*SKP2* coding sequence into mouse *Pbsn* locus, such that hSKP2 expresses from the modified *Pbsn* locus. Because *Pbsn* gene is X-linked, a solution to maintain transgenic line is to crossbreed of heterozygous females with wild type males. Since the CRISPR/Cas9 approach more accurately targets of hSKP2 to endogenous mouse *Pbsn* locus, the time and effort for screening of founder animals is significantly reduced compared to random insertion approaches. Indeed, our approach results in a significantly high rate (about 44.2%) of positive transgenic lines of h*SKP2*-KI and is more efficient compared to traditional transgenic approaches. In addition, endogenous mouse *Pbsn* promoter driven hSKP2 overexpression would be more physiopathologically relevant and provide more reproducible phenotypes and molecular signatures compared to traditional random insertion approaches into an unknown locus(s). In our study, hSKP2 is specifically overexpressed in all lobes of mouse prostate, but not in other organs, such as testis, kidney, spleen, and liver. SKP2 overexpression in mouse prostates induces atypical cell proliferation, leading to a progressive development of spontaneous prostatic lesions including hyperplasia, mPIN, and low-grade prostatic adenocarcinoma over 14 months of age.

Compared to the ARR2PB (the rat *probasin*) promoter driven hSKP2 transgenic mice [40], our probasin-h*SKP2*-KI mice similarly develop hyperplasia, low grade PIN, high grade PIN and low-grade carcinoma, but exhibit enlarged prostate glands and low-grade carcinoma in a slower fashion at the advanced age of 14 months. The downregulation of p27^Kip1^, a key substrate of SKP2 was also observed to be restricted in the hyperplastic and dysplastic regions, accompanied by SKP2 overexpression and a marked increase in Ki67 positively staining cells.

The previously unreported discovery in this study stems from the RNAseq analysis, which have identified the top enrich pathways, including EMT, collagen fibril organization, interferon α/γ pathways, in prostate tissues from the h*SKP2* KI vs. wild-type mice. Deconvolution of bulk RNA seq data into single cell distribution shows a significant increase in fibroblast population and proliferative classic monocytes and a decrease in CD8^+^ T cells, B cells, and myeloid dendritic type I cells in the prostate of the h*SKP2* KI mouse. Ruan et al. [41] reported that SKP2 stabilized the TWIST1 protein, a master regulator of EMT, through a non-degradative ubiquitination to promote EMT and acquisition of cancer stem cell properties. We also show that SKP2 overexpression in prostate cancer PC3 cells increases the protein levels of TWIST1 and the expression of EMT related genes, including *FMOD, THY1, NFIL3* and *IRF2*, leading to a marked increase in migration and invasion. All these results are in alignment with the fact that SKP2 overexpression or amplification occurs more frequently in prostate cancer tissues with Gleason scores of 9 or 10 and in mesenchymal tumors, like sarcoma. You et al. reported that TWIST1 regulated the basal expression levels of interferon regulatory factor 9 (IRF9) and interacted with IRF9 to promote IRF9 binding to the IFN-stimulated response element [42].

Other studies also showed that SKP2 interacts with IFN-I receptor 2, T-box expressed in T cells, and SH2-containing protein tyrosine phosphatase-2 to regulate IFN-a/γ abundance [43, 44]. This emerging evidence has pointed to a central role of SKP2 in regulating IFN-a/γ in the tumor microenvironment through multiple mechanisms. Therefore, targeting SKP2 in both cancer cells and tumor microenvironment would be a robust strategy for prostate cancer prevention and treatment.

To facilitate the efficient screening of novel selective SKP2 inhibitors, we have further developed paired prostate organoid cultures from both the h*SKP2*-KI and wild-type mice. While both the C1 inhibitor and the SKP2 degrader FKA selectively inhibit the growth of SKP2 overexpressing prostate organoids over normal prostate organoids at the early stage of the treatments, the SKP2 inhibitor C1 that targets SKP2/Cks1 interaction lost its activity against the growth of SKP2 overexpressing prostate organoids after 8 days of its treatments. However, FKA by degrading SKP2 protein continuously inhibits the growth of the SKP2 overexpressing prostate organoids over time. Currently, there are no effective or suitable SKP2 inhibitors for clinical trials yet. Our organoid cultures would provide a novel tool to screen and test new more potent SKP2 inhibitors for pre-clinical and clinical studies, and to understand potential mechanisms of resistance to SKP2 inhibitors with different targeting mechanisms.

In summary, the prostate-specific overexpression of h*SKP2* in mice initiates spontaneous carcinogenesis in a time-dependent manner and leads to a significant remodeling of the prostatic microenvironment through inducing EMT, increasing fibroblast population and enhancing extracellular matrix accumulation. These changes are in parallel with down-regulated IFN a/γ and anti-tumor immunity signaling, enabling the evasion of immune surveillance to facilitate the process of prostate tumorigenesis. Our prostate-specific SKP2 humanized mouse model offers a valuable platform for investigating mechanisms of immune evasion and tumor microenvironment reprogramming during prostate cancer initiation and progression, exploring the combined effects of oncogenes and tumor suppressors, and evaluating potential agents targeting human SKP2 for prostate cancer prevention and therapy.

## Acknowledgment

This work was supported in part by NIHUG3 CA290368, NIHR01 CA260351-02, and VA merit award I01BX005105 (to X. Zi.). In addition, the authors acknowledge support from the Chao Family Comprehensive Cancer Center (NCI award P30CA62203) and its shared resources.

## Authors’ Contributions

**L. Song:** Data curation, formal analysis, investigation, methodology, writing–original draft, reviewing and editing. **V. Nguyen:** Data curation. **S. Xu:** Data curation. **K. Ho:** Data curation. **BH. Hoang:** Resources. **E. Uchio:** Resources. **J Yu:** Resources. **X. Zi:** Conceptualization, resources, data curation, formal analysis, supervision, funding acquisition, validation, investigation, visualization, methodology, writing–original draft, project administration, writing–review and editing.

