## Supplemental Figures and Table for "The Establishment of Prostate-specific, SKP2 Humanized Mice by CRISPR Knock-in Method Reveals Neoplastic Initiation and Microenvironmental Reprogramming"

**Supplementary Figures and Table**

**Supplementary Figure 1. A.** Quantitative PCR analysis of the expression of interferon-related genes *P2ry14*, *Usp18* and *Ifitm3* in the prostate of h*SKP2*-KI mice. N = 4, **P*<0.05. B. Quantitative PCR analysis of the expression of interferon-related genes *PFKP*, *USP18* and *LATS2* in human prostate cancer PC3 cells overexpressing SKP2. N = 3, **P*<0.05, ****P*<0.001.

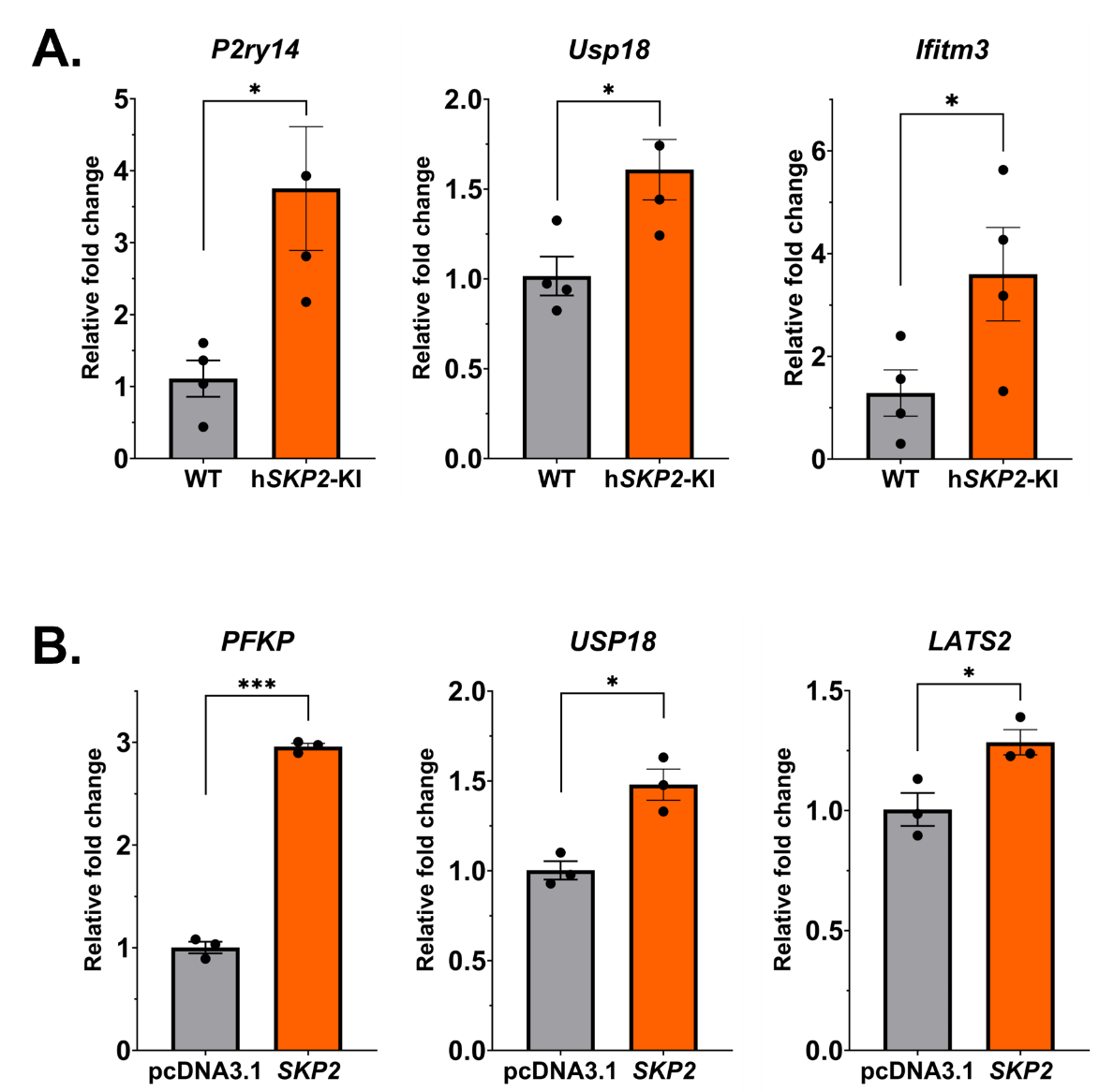

**Supplementary Figure 2.** **A**. WT and h*SKP2*-KI mouse prostate organoids were treated with FKA at 12.5µM for 10 days. Viabilities of these organoids were examined by adding a fluorescent dye CalAM for 30 minutes before fluorescent intensity reading and imaging at indicated time points, Scale bar: 100 µm. **B**. Relative fluorescent intensities of these organoids to their baseline levels at day 0 before treatments were calculated at indicated time points and presented as a line graph.

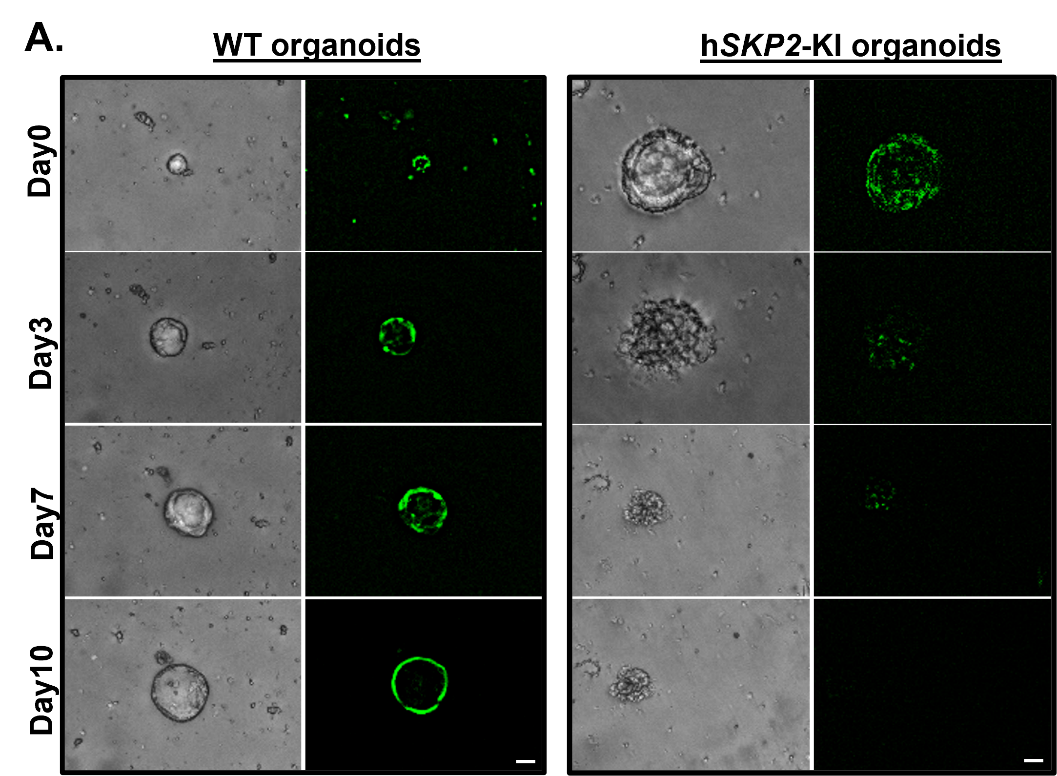

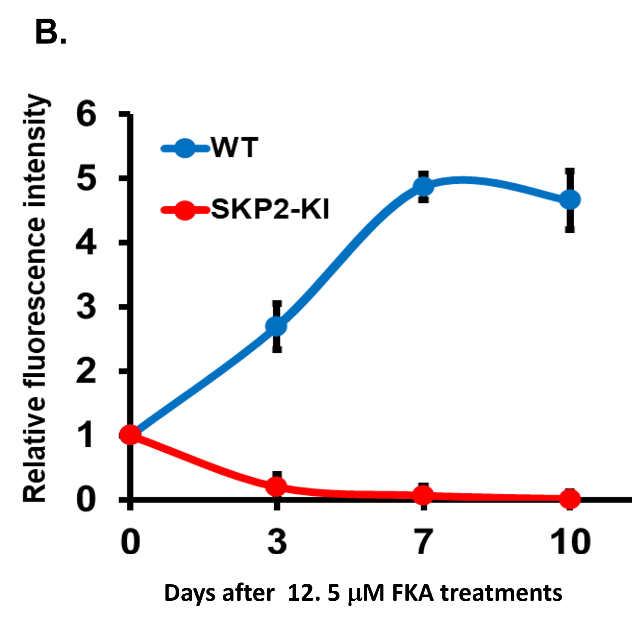

**Supplementary table 1.** Quantitative PCR primers

| **Gene** | **Species** | **Forward sequence (5'-3')** | **Reverse sequence (5'-3')** |
| --- | --- | --- | --- |
| *Fmod* | mouse | AGCAGTCCACCTACTACGACC | CAGTCGCATTCTTGGGGACA |
| *Thy1* | mouse | TGCTCTCAGTCTTGCAGGTG | TGGATGGAGTTATCCTTGGTGTT |
| *Wnt5a* | mouse | CAACTGGCAGGACTTTCTCAA | CATCTCCGATGCCGGAACT |
| *Cd38* | mouse | GGTCCAAGTGATGCTCAATGGG | AGCTCCTTCGATGTCGTGCATC |
| *Usp18* | mouse | GGAACCTGACTAAGGACCAGATC | GAGAGTGTGAGCAGTTTGCTCC |
| *Il15* | mouse | GTAGGTCTCCCTAAAACAGAGGC | TCCAGGAGAAAGCAGTTCATTGC |
| *Pfkp* | mouse | AAGAGGAAACCAAGCAGTGCGC | TTCCTCGGAGTTTCACGGCTTC |
| *Lats2* | mouse | GCACTGGATTCAGGTGGACTCA | CGACAGTTGGAAACATCGTCCC |
| *P2ry14* | mouse | CGACAGTTGGAAACATCGTCCC | GCTGTAGTGACCTTCCGTCTGA |
| *Tnfaip2* | mouse | TGTGCACCTGCACCTAGTGAA | CACTGGAATCTTGCAGGCGAA |
| *Gapdh* | mouse | CTCCCACTCTTCCACCTT CG | GCCTCTCTTGCTCAGTGTCC |
| *Ifitm3* | mouse | CCCCCAAACTACGAAAGAATCA | ACCATCTTCCGATCCCTAGAC |
| *SKP2* | human | CTGAGCTGCTAAAGGTCTCTGGTG | CACCCCTTGAGACAGCAACCGAC |
| *FMOD* | human | ATTGGTGGTTCCACTACCTCC | GGTAAGGCTCGTAGGTCTCATA |
| *THY1* | human | ATGAAGGTCCTCTACTTATCCGC | GCACTGTGACGTTCTGGGA |
| *WNT5a* | human | GCCAGTATCAATTCCGACATCG | TCACCGCGTATGTGAAGGC |
| *IFITM3* | human | CTGGGCTTCATAGCATTCGCCT | AGATGTTCAGGCACTTGGCGGT |
| *ACTB* | human | CACCATTGGCAATGAGCGGTTC | AGGTCTTTGCGGATGTCCACGT |
| *TNFAIP2* | human | TGCTCCAGAACCTGCATGAGGA | AACTCAGGCAGCCTCGTGTCTA |
| *USP18* | human | TGGACAGACCTGCTGCCTTAAC | CTGTCCTGCATCTTCTCCAGCA |
| *IL15* | human | AACAGAAGCCAACTGGGTGAATG | CTCCAAGAGAAAGCACTTCATTGC |
| *PFKP* | human | AGGCAGTCATCGCCTTGCTAGA | ATCGCCTTCTGCACATCCTGAG |
| *LATS2* | human | GTTCTTCATGGAGCAGCACGTG | CTGGTAGAGGATCTTCCGCATC |
| *P2RY14* | human | GCCGCAACATATTCAGCATCGTG | GCTGTAATGAGCTTCGGTCTGAC |
